# Discovering dynamical models of human behavior

**DOI:** 10.1101/2022.03.20.484666

**Authors:** Paul I. Jaffe, Russell A. Poldrack, Robert J. Schafer, Patrick G. Bissett

**Author notes:** These authors contributed equally to this work (Russell A. Poldrack, Robert J. Schafer, Patrick G. Bissett).

## Abstract

Response time (RT) data collected from cognitive tasks are a cornerstone of psychology and neuroscience research, yet existing models of these data either make strong assumptions about the data generating process or are limited to modeling single trials. We introduce task-DyVA, a deep learning framework in which expressive dynamical systems are trained to reproduce sequences of RTs observed in data from individual human subjects. Models fitted to a large task-switching dataset captured subject-specific behavioral differences with high temporal precision, including task-switching costs. Through perturbation experiments and analyses of the models’ latent dynamics, we find support for a rational account of switch costs in terms of a stability-flexibility tradeoff. Thus, our framework can be used to discover interpretable cognitive theories that explain how the brain dynamically gives rise to behavior.

## Introduction

Developing dynamical models that capture how people integrate information and make decisions in real time is a fundamental problem in psychology and neuroscience. Cognitive psychologists study decision-making and other behaviors using cognitive tasks in controlled laboratory settings, and RT data collected from these tasks contain rich and complicated temporal dependencies that inform models of the underlying mental phenomena (*1, 2*). There is a long and fruitful tradition in psychology of explaining these data with cognitive process models, in which simple and interpretable components are assembled in a “top-down” fashion so as to capture behavioral effects of interest (*3, 4*). However, the handcrafted nature of these models entails a set of strong assumptions that may be difficult or impossible to verify. More recently, deep neural network models and recurrent neural network (RNN) models trained to perform cognitive tasks have emerged as a complementary modeling paradigm (*5–7*). Like their connectionist predecessors (*8, 9*), modern neural network models impose a more limited set of structural assumptions, allowing the constraints of the task to produce behavior emergently. However, these models have not been extended to capture RT data spanning multiple trials, a significant departure from realistic human behavior. Moreover, the vast majority of these models as they have been applied in neuroscience and psychology are trained to perform cognitive tasks—rather than fit with behavioral data (c.f. (*10, 11*))—and thus cannot be used to model individual differences.

To address these limitations, we introduce task-DyVA (“dee-vuh”), a novel framework for modeling sequential RT data grounded in a recently proposed class of machine learning models, dynamical variational autoencoders (*12*). Our approach combines the expressive power of RNNs with the ability to capture individual differences in behavior: each task-DyVA model is directly constrained to reproduce sequences of RTs observed in data from a single human participant. In brief, our approach furnishes generative, dynamical models that simulate how humans perform cognitive tasks in real time.

We apply our framework to investigate task-switching, a well-established experimental paradigm used to study fundamental aspects of cognitive control and mental flexibility (*13–16*). In task-switching experiments, subjects are cued to perform one of two or more discrete tasks on each trial. For example, in the task switching game “Ebb and Flow” (Lumosity) studied here (*16*), subjects are shown a set of moving leaves on each trial and must report either their direction of motion or their orientation (Fig. 1A). The appropriate type of response (direction or motion) is cued by the color of the stimuli. While task-switching experiments vary considerably in methodology, a consistent empirical observation is that subjects are slower to respond and less accurate on trials in which the task switches (“switch” trials) relative to trials in which the task remains the same (“stay” trials) (*14, 15*). This switch cost provides a strong test of the ability of task-DyVA to capture history-dependent behavioral phenomena: in order to generate a switch cost, the state of the model’s latent dynamics in the present must contain information about the stimuli experienced in the past.

**Fig. 1:**
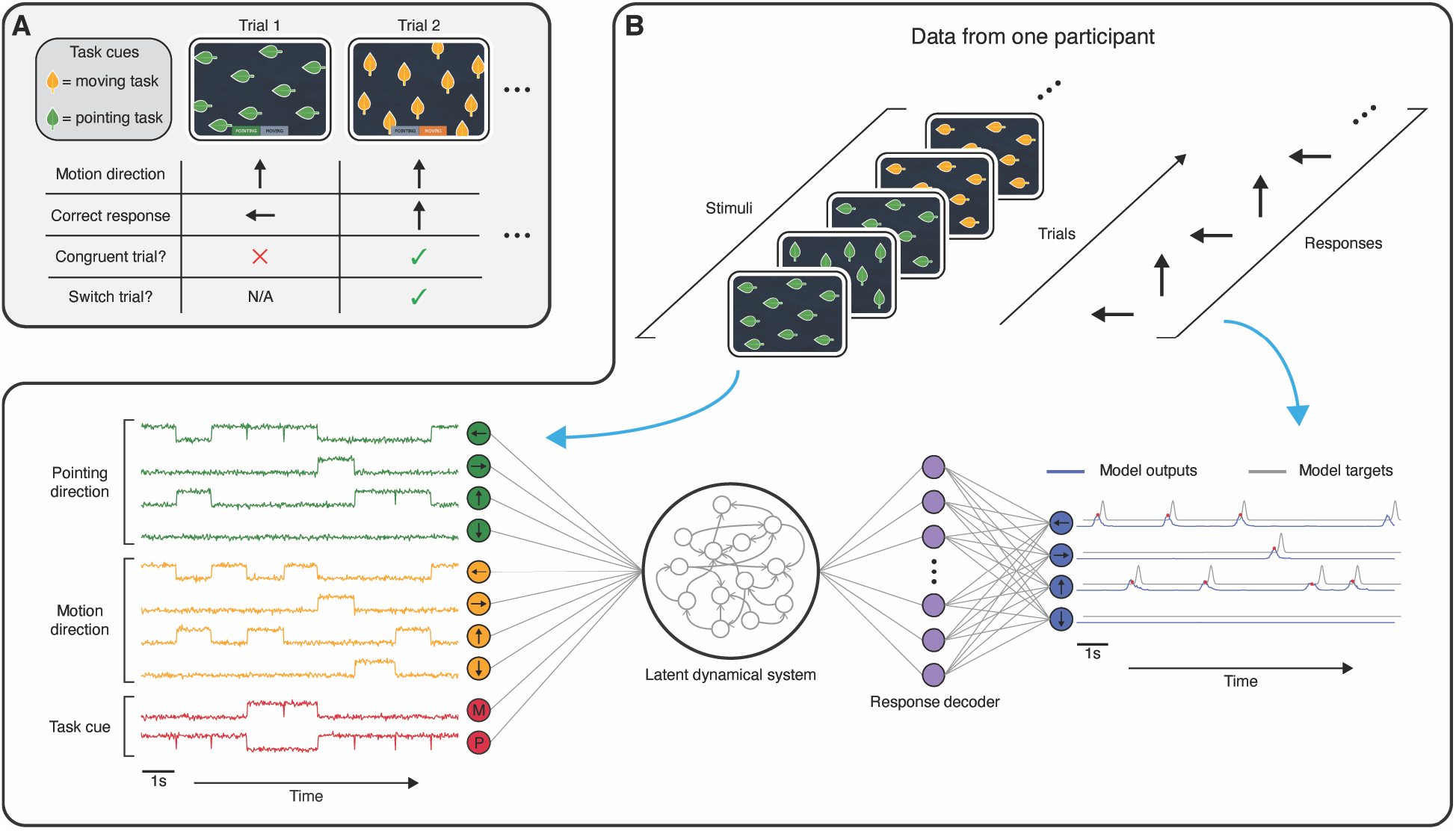
Task-DyVA modeling framework and task-switching game. (**A**) Ebb and Flow, a task-switching game (Lumosity). Congruent trial: moving and pointing stimuli are oriented in the same direction. Switch trial: task on trial *n* - 1 was different from task on trial *n*. (**B**) Schematic of task-DyVA. Top: Ebb and Flow gameplay data from one participant. Bottom left: the transformed stimuli supplied as inputs to the model. Bottom right: Model outputs (blue) and model output targets (gray). Red dots indicate the model’s responses.

Using a large gameplay dataset from the Lumosity cognitive training platform, we fit subject-specific models to data from the task-switching game Ebb and Flow. The fitted models reproduced a range of behavioral phenomena with high temporal precision, including the mean RT, stimulus congruency effects, and switch costs. Adopting a state space perspective, we discovered that the two tasks were encoded in different regions of the model’s latent space and that the switch cost was directly related to the transit time between these task regions. Moreover, the separation of task spaces conferred robustness to noise. Our findings therefore provide a computational basis for a normative explanation of switch costs originally proposed by Musslick & Cohen (*17*): switch costs emerge *because* separable task representations confer behavioral stability.

### The task-DyVA modeling framework

The fundamental operating principle of task-DyVA is simple: we train an expressive dynamical system to take experimentally observed sequences of task stimuli as inputs and generate observed sequences of task responses as outputs (Figs. 1B and S1). The product of this training is a generative model that can be used to simulate sequential human behavior on cognitive tasks, and whose latent dynamics can be flexibly queried to gain insight into the model’s representation of the task. At the heart of the generative model is a latent dynamical system with state variables **z**_*t*_ that is intended to capture all of the internal cognitive operations required to perform the task. The latent state evolves according to:

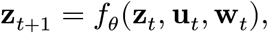

where **u**_*t*_ is the task stimulus at timestep *t*, **w**_*t*_ is a noise term, and *f*_*θ*_ is the dynamics function. Similar to RNNs (*18*), *f*_*θ*_ is highly expressive: it is capable of approximating a large family of dynamical systems (see supplementary materials). This formulation imposes few constraints on the nature of the dynamics to be recovered, effectively allowing the data rather than our prior assumptions to shape the learned task representation.

Each stimulus modality × direction combination is represented with a binary-valued unit indicating its presence or absence at each moment in time (Fig. 1B, bottom left). Task cues are represented in a similar fashion. Importantly, whether a given trial is a switch versus a stay trial is not explicitly coded in the model input and consequently must be learned from the temporal structure of the stimuli. To facilitate model training and to increase biological plausibility, zero-mean Gaussian noise (0.1 SD) is added independently to each stimulus unit.

Model outputs **x**_*t*_ are generated by the decoder model *p*_*θ*_(**x**_*t*_|**z**_*t*_), parameterized by a multilayer perceptron (MLP). Each possible task response direction is represented by a separate output channel. The activation of a given output channel at any moment in time is related to the probability that the model will generate a response in that direction (see supplementary materials). To train the model, we require an error term that measures how close the model outputs are to the participant’s responses. This is accomplished by centering a Gaussian kernel at each RT (SD = 50ms), forming a smooth response template (Fig. 1B, bottom right). To learn the parameters of the generative model, we employ an approximate inference framework known as the variational autoencoder or VAE (*12, 19, 20*) (see supplementary materials).

In sum, the generative model defined above obeys state space assumptions: given **u**_*t*_ and **w**_*t*_, the current response **x**_*t*_ and future state **z**_*t*+1_ depend only on the current state **z**_*t*_ (see Fig. S1A for a probabilistic graphical model). The state space formulation of our model is important because it aids interpretability: all of the information pertaining to the progression of computation in the task can be observed directly from the latent state **z**_*t*_ at each moment in time.

## Results

### Task-DyVA emulates human task-switching behavior

We fit a separate task-DyVA model to Ebb and Flow data from each of 140 participants (20 participants in each of seven decade-long age bins, ages 20 to 89). To reduce response variability from early stages of task learning, we selected participants who had practiced extensively and used gameplays from a late stage of practice for model training (gameplays 150 to 500, inclusive). After transforming the data as described above, each gameplay was segmented into short sequences for model training (5s duration, ∼3 to 4 trials). After training, we used the fitted models to generate responses on longer stimulus sequences from a holdout dataset (10s duration, ∼6 to 8 trials). The model’s responses were compared to the participant’s actual responses using the same set of stimuli (Figs. 2A-2C and S2).

**Fig. 2:**
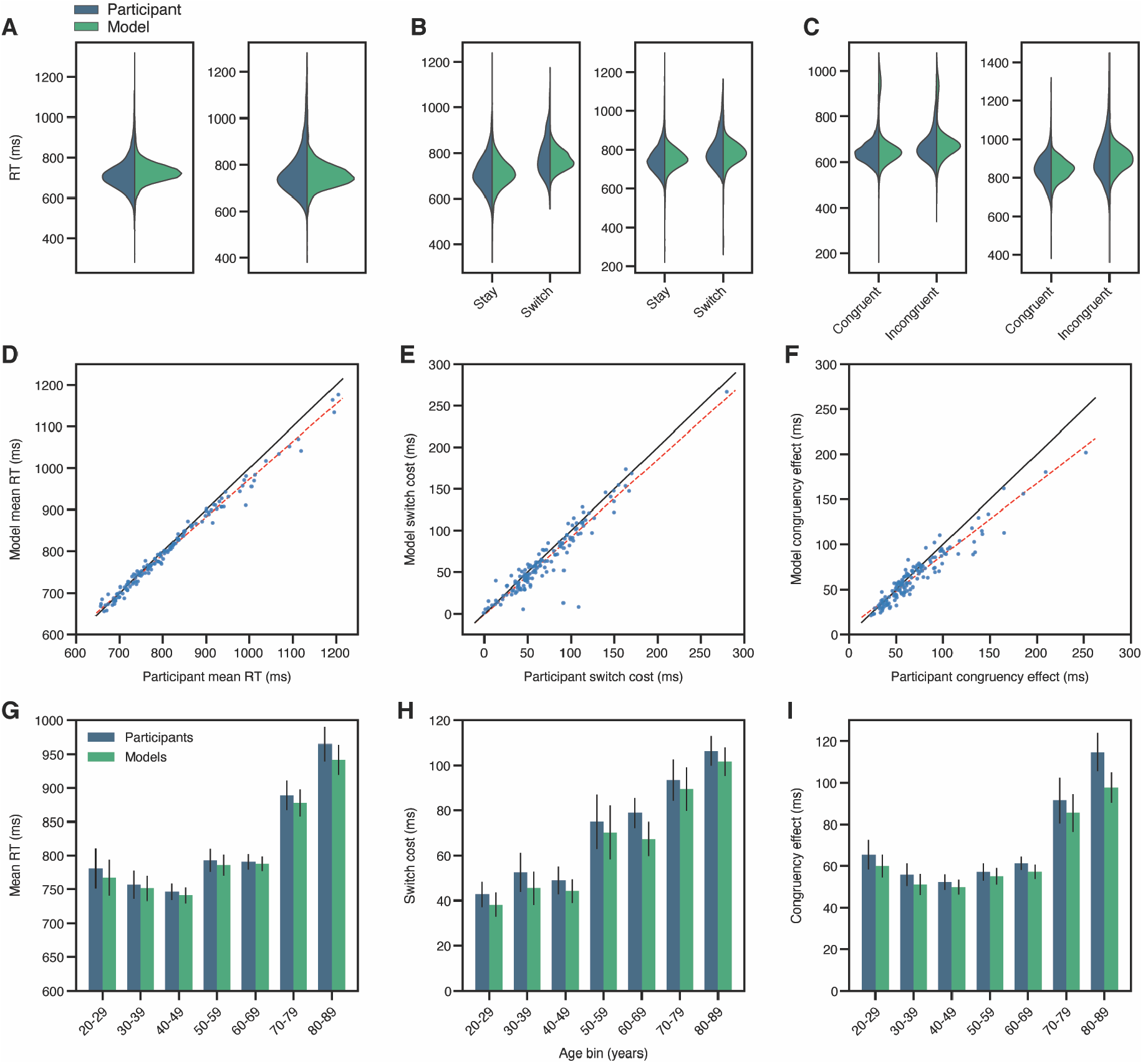
Task-DyVA models capture human behavior. (**A**) Example RT distributions for two participants and corresponding fitted models. (**B**) Example stay and switch trial RT distributions for two participants/models. (**C**) Example congruent and incongruent trial RT distributions for two participants/models. (**D**) Mean RTs. (**E**) Switch costs. (**F**) Congruency effects. For panels D-F, each point is one participant/model; black line: unity; red dashed line: best linear fit; N = 140 participants/models. (**G**) Mean ± s.e.m. mean RTs within each age bin (N = 20 participants/models per age bin). (**H)** Mean ± s.e.m. switch costs. (**I**) Mean ± s.e.m. congruency effects.

We assessed the performance of each model on three key behavioral metrics: the mean RT, the switch cost (defined as the difference in mean RT on switch trials versus stay trials, and the congruency effect (defined as the difference in mean RT on incongruent versus congruent trials; see Fig. 1A). For each of these metrics, the responses generated by the models were highly correlated with those of the participants and the slopes of a linear best-fit line were close to but slightly less than one (Figs. 2D-2F; mean RT Pearson’s *r* = 0.99, test for non-zero correlation using the exact distribution of *r:* p < 1e-134, best-fit slope = 0.91; switch cost Pearson’s *r* = 0.94, p < 1e-66, best-fit slope = 0.93; congruency effect Pearson’s *r* = 0.96, p < 1e-74, best-fit slope = 0.80; N = 140 participants/models). The fitted models also captured age-dependent trends in each of these behavioral metrics (*16*) (Figs. 2G-2I).

At the level of stimulus noise used to train the models (0.1SD), the trained models typically exhibited higher accuracy (∼99%) and less RT variability (∼74ms SD) relative to their corresponding participants (Figs. S3 and S4; participant accuracy: ∼96%, participant RT SD: ∼124ms). As expected, increasing the magnitude of the stimulus noise reduced model accuracy (Fig. S3A). More notably, increasing the stimulus noise also revealed differential effects on stay versus switch trial accuracy and congruent versus incongruent trial accuracy, effects also observed in the participants and in other task-switching studies (*14, 15*) (Figs. S3B-S3C). At 0.4SD noise, the magnitudes of the ‘accuracy switch cost’ and ‘accuracy congruency effect’ of the trained models were not significantly different from those of the participants (p = 0.97 and p = 0.14, respectively, signed-rank test). The correlations between participant and model mean RTs, congruency effects, and switch costs were reduced at this elevated noise level, but remained highly significant (Fig. S5). Since our primary goal was to leverage the dynamic nature of task-DyVA to understand the learned representation of the RT effects (mean RT, RT switch cost, and RT congruency effect), we elected to fix the stimulus noise to the original magnitude in the remainder of our analyses (0.1SD), for which the fit between participant and model RT effects was best.

### A hierarchical representation of the task

The close correspondence between participant and model behavior validates task-DyVA as a tool that can be used to discover how humans dynamically represent cognitive tasks. Accordingly, we investigated the models’ representation of Ebb and Flow by examining the latent dynamics in relation to the task cues and stimuli. We separated trials according to the active task cue and task-relevant stimulus direction and calculated the trial-averaged latent state trajectories aligned to stimulus onset for each group of trials. To aid visualization, we projected these trajectories onto their top three principal components (PCs) which accounted for the vast majority of variance (∼91%) in the latent state (Figs. 3A-3B, and S6; variance calculated across all trials and timepoints). In the example model shown in Fig. 3A, the smooth curves correspond to trial-averaged trajectories of the latent state for a given trial type. For example, the solid blue line corresponds to all trials in which the task cue was the moving task and the stimuli were moving left.

**Fig. 3:**
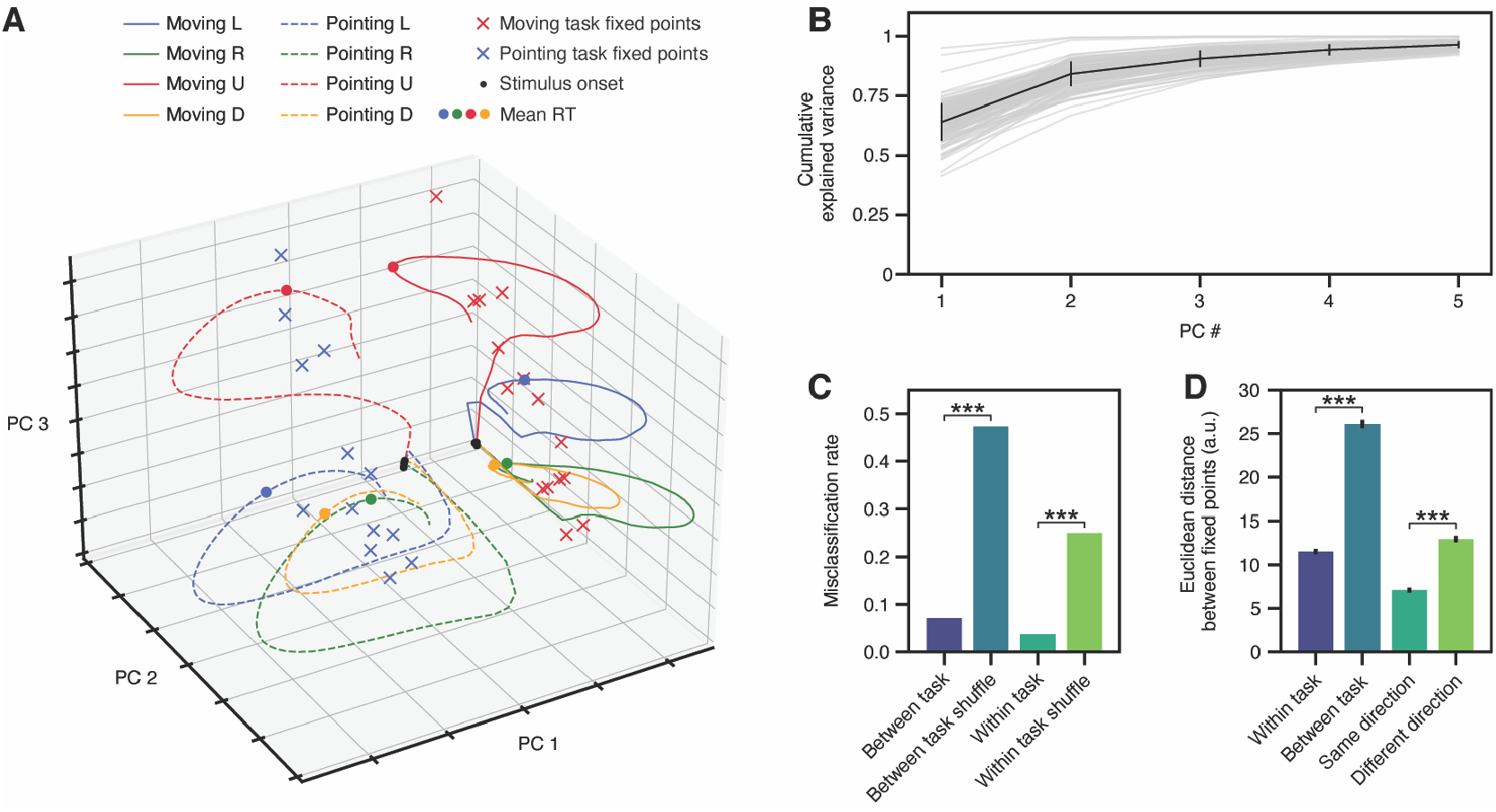
The latent representation of the task is organized hierarchically. (**A**) Trial-averaged latent state trajectories for each task-relevant stimulus direction × task cue combination and stable fixed points for one model. (**B**) Cumulative explained variance vs. PC number (gray lines: individual models; black line: mean ± SD). (**C**) Mean misclassification rate of the LDA models used to classify latent state vectors. Error bars were too small to visualize (s.e.m. < 1e-3 for all conditions). (**D**) Euclidean distance between pairs of stable fixed points (mean ± s.e.m.). ***p < 1e-23, signed-rank test.

The latent state trajectories shown in Fig. 3A exemplify a notable feature of the learned task representations: the two tasks are represented in different regions of the latent space (compare solid and dashed trajectories; see also Fig. S6). To quantify this separation for each model, we trained a linear discriminant analysis (LDA) model to classify latent state vectors from each trial according to which task cue was active (Fig. 3C, two leftmost bars). The latent state vectors were derived from onset-aligned latent state trajectories evaluated at the model’s mean RT. The mean error rate of these LDA models was ∼7%, significantly less than the error rate of LDA models trained on shuffled data (∼47%; p < 1e-23, signed-rank test). Thus, the latent state vectors could be reliably assigned to one or the other task, indicating that the representations of the two tasks were well-separated in the latent space. For trials corresponding to a given task cue, we also used LDA to quantify how well the latent state vectors from trials with a given task-relevant stimulus direction could be distinguished from those of the other three stimulus directions (Fig. 3C, two rightmost bars). The mean error rate of these LDA models was ∼4%—significantly less than the error rate for shuffled data (∼25%, p < 1e-23, signed-rank test)—indicating that the correct response direction was readily determined from the position of the latent state.

To understand the dynamics of the learned task representations, we identified a set of stable fixed points for each model corresponding to points in the latent space that absorb nearby trajectories (i.e., attractors) (*5, 21*). Fixed points were identified by running the models in ‘generative mode’ using long (50s) sequences of static stimuli as inputs; each of the 32 possible stimulus configurations was used to discover fixed points (see supplementary materials).

Fixed points identified with a given task cue were localized to a task-specific region of the latent space, mirroring the separation of latent state trajectories that we observed (see ‘x’ marks in Figs. 3A and S6). To quantify this separation, we bucketed pairs of fixed points according to whether or not they were identified with the same task cue and calculated the mean Euclidean distance between all pairs of fixed points within each bucket (Fig. 3D, two leftmost bars). Pairs of fixed points identified with the same task cue were indeed closer together than pairs identified with different task cues (p < 1e-23, signed-rank test, N = 137 models). For a given task cue, we also found that pairs of fixed points corresponding to the same task-relevant stimulus direction were closer together on average than those corresponding to different task-relevant stimulus directions (Fig. 3D, two rightmost bars; p < 1e-23, signed-rank test, N = 137 models). Thus, the fixed point landscape of the trained models revealed that the task was represented in a hierarchical fashion. Globally, each task was associated with a circumscribed attractor region that funneled the latent state according to the active task cue. Locally, within each task region, the latent state was pulled toward fixed points corresponding to the correct response direction.

### Dynamical origins of the switch cost

One consequence of the hierarchical representation we observed is that the representations for the two tasks are well-separated within the latent space. Conceivably, this separation contributes to switch costs, simply because it takes time for the latent state to transition from one task region to the other (Fig. S7A). However, two issues complicate this simple interpretation: it is not necessarily costly to move large distances in the latent space and, more generally, the time it takes for the latent state to travel between two points along a trajectory is determined by the model’s dynamics (i.e., the vector field), not Euclidean distance. Thus, in principle, the latent state could transition from task region A to B relatively quickly on switch trials, with switch costs resulting primarily from slower dynamics within task region B (Fig. S7B).

Despite this complexity, the hypothesis that greater separation between task regions contributes to switch costs makes two predictions that were borne out by the models (see examples in Figs. 4A and S7C). First, if the transition between task regions A and B is costly (i.e. slow), then we should observe a positive correlation between the distance to task region B at stimulus onset and RTs for switch trials (Fig. S7A). We assessed this by calculating the Euclidean distance between the latent state at stimulus onset and a single task-specific reference point for each task, points referred to as task centroids (see supplementary materials). We found that the vast majority of models exhibited a positive correlation between RTs and distance on switch trials (Fig. 4B; number of models with positive Pearson’s *r*: 134/140; mean ± s.e.m. *r*: 0.24 ± 0.012; sign-rank test for non-zero population *r*: p < 1e-23). A second prediction of this hypothesis is that switch costs should be positively correlated with the distance between task centroids across the models for different individuals. We measured the distance between task centroids using a normalized distance measure that accounts for potential differences in the overall scale of different models (see supplementary materials). We observed a strong correlation between switch costs and the distance between task centroids (Fig. 4C; Pearson’s *r* = 0.65; test for non-zero correlation using the exact distribution of *r*: p < 1e-17, N = 140 models).

**Fig. 4:**
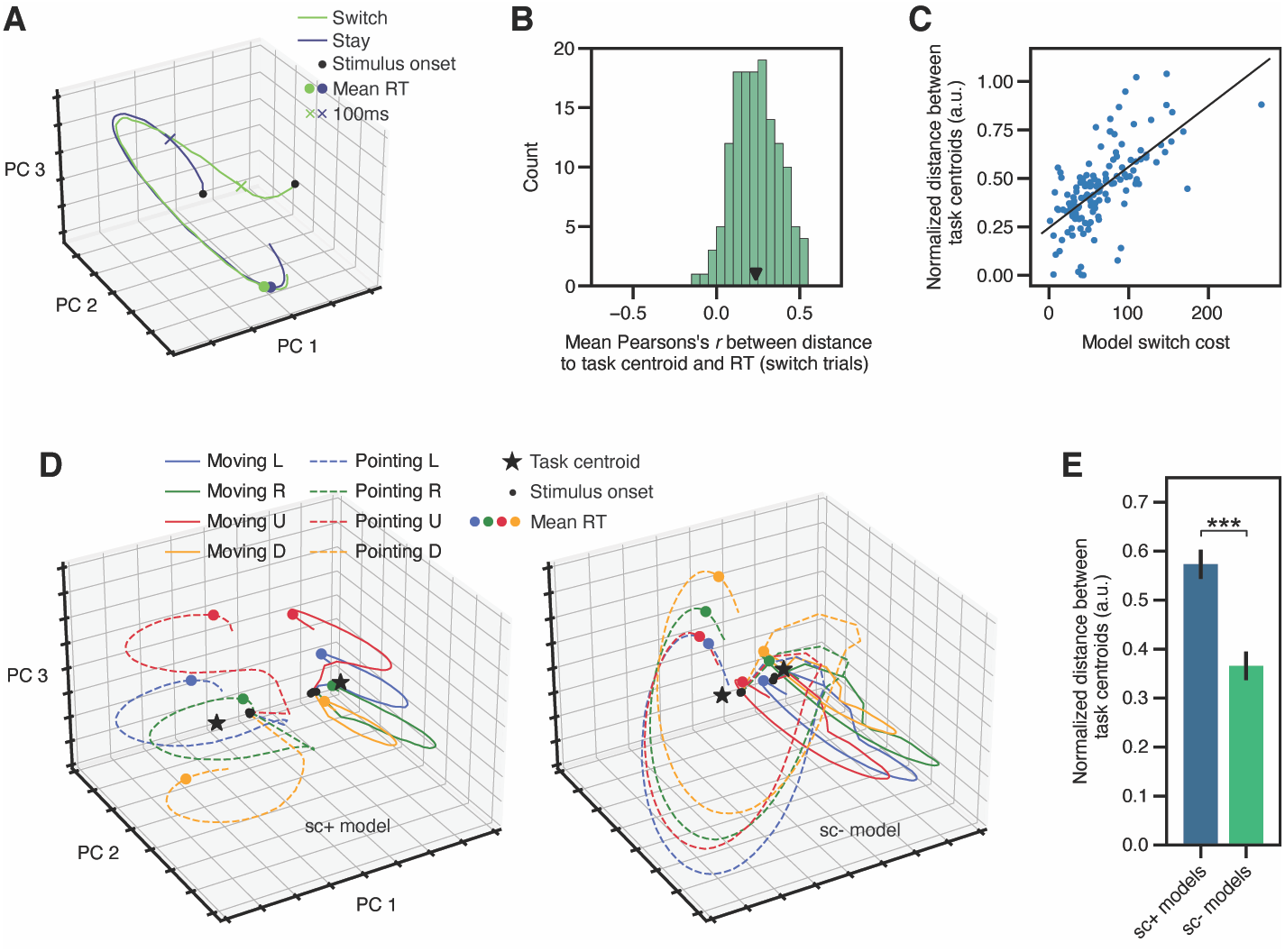
Separated task representations contribute to switch costs. (**A**) Latent state trajectories from one model averaged over trials with a fixed stimulus configuration (same stimuli and task cues on the current trial; task cues on the previous trial differed to select stay vs. switch trials). (**B**) Mean Pearson’s *r* between the Euclidean distance to the task centroid at trial onset and RT on switch trials (sc+ models, N = 140; black triangle shows the population mean). For each model, *r* was calculated separately for trials with a fixed stimulus configuration, then averaged across stimulus configurations. (**C**) Model switch cost vs. the normalized distance between task centroids (sc+ models, N = 140; black line shows the best linear fit). (**D**) Trial-averaged latent state trajectories for sc+ (left) and sc- (right) models trained on data from the same participant. The sc+ and sc- plots have the same axis limits. (**E**) Mean ± s.e.m. normalized distance between task centroids (N = 25 sc+ models and 25 sc- models). ***p < 1e-4, signed-rank test.

As a more direct test of the idea that the separation between task regions contributed to switch costs, we leveraged a unique strength of task-DyVA: the models can be trained with synthetic behavioral data in which features of interest have been removed (or augmented). In particular, we reasoned that trained models that do not exhibit a switch cost would also exhibit reduced separation of the latent task representations. We set out to create such a scenario by training models using synthetic data lacking a switch cost. To this end, we selected 25 participants with large switch costs from the original cohort of 140 participants/models and modified the data from these participants to remove the switch cost (see supplementary materials). We trained a task-DyVA model on each of these modified datasets, allowing us to compare the two models trained with the same participant’s data (which either did or did not have a switch cost). To the extent possible, all other aspects of the training data, model architecture, and training procedure did not differ from the original models. For brevity, we refer to the models trained on synthetic data lacking a switch cost as the sc-models, and the original models trained on data with a switch cost as the sc+ models.

As expected, the switch cost of the sc- models was greatly reduced relative to that of the sc+ models (Fig. S8A; mean ± s.e.m. switch cost for sc+ models: 111.4 ± 4.7ms, for sc- models: 5.4 ± 1.4ms, p = 1.2e-5, signed-rank test, N = 25 models). In contrast, the mean RT and congruency effect of the sc- models differed only marginally from the sc+ models (Figs. S8B-S8C). Relative to the sc+ models, the distance between task centroids in the sc-models was significantly reduced (Figs. 4D-4E; p = 8.1e-5, signed-rank test). Thus, the distance between task regions—a consequence of the hierarchical task representation—contributed to larger switch costs both within and across models.

### The separation of task representations confers robustness

We next examined whether the observed structure of the latent representation conferred any functional benefits, beyond providing a computational explanation for switch costs. In particular, we considered the possibility that the increased separation of task regions in the sc+ models would confer robustness, e.g. by reducing the likelihood that noise would push the latent state into an incorrect response region. We leveraged the reduced task separation that we observed in the sc- models to test this idea, varying the magnitude of the stimulus noise and assessing the effect on model accuracy for the sc+ and sc- models. Relative to the sc-models, the accuracy of the sc+ models was less affected by stimulus noise for noise magnitudes >= 0.3 SD (Figs. 5A-B and S9A; p < 0.05 for all noise values between 0.03 and 1, inclusive, signed-rank test, N = 25 models).

**Fig. 5:**
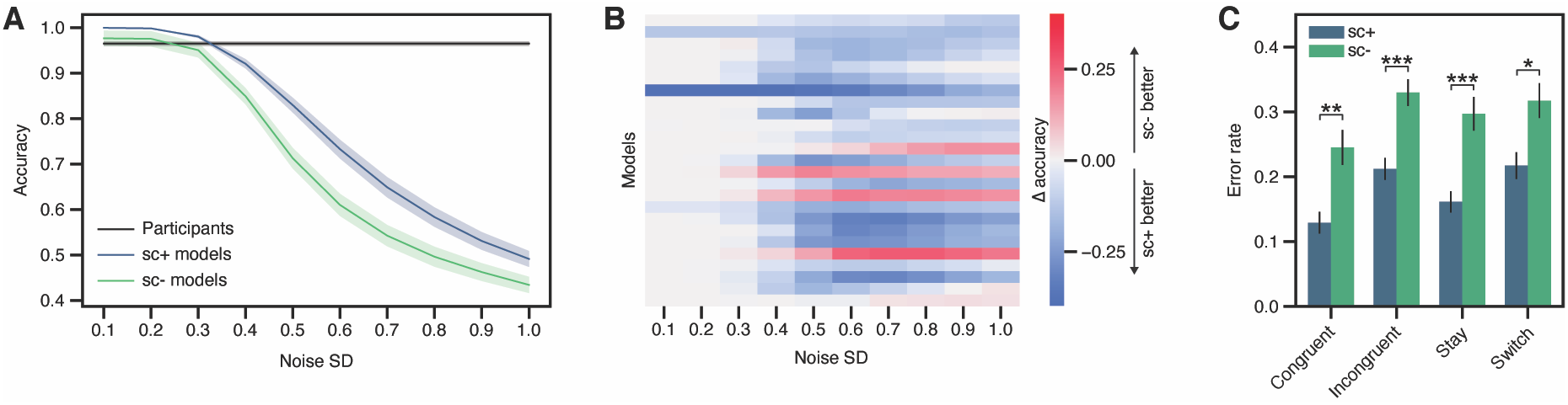
Models with more separated representations are more robust to noise. (**A**) Mean ± s.e.m. model accuracy vs. stimulus noise magnitude (sc+: models trained on data that has a switch cost; sc-: models trained on data that lacks a switch cost; N = 25 models). Note that participant accuracy was not assessed at different noise levels. (**B**) Difference in accuracy between all pairs of sc+ and sc- models. Blue colors indicate that the sc+ model has higher accuracy than the paired sc- model. (**C**) Error rate conditioned on trial type (mean ± s.e.m.; noise SD = 0.5). *p < 0.05, **p < 0.01, ***p < 0.001, signed-rank test.

We next tested whether the reduced task separation in the sc- models would predispose them to a different pattern of errors versus the sc+ models. To assess this, we calculated the conditional error rate for four different trial types: congruent, incongruent, stay, and switch (Figs. 5C and S9B-E). For all four trial types, the error rate was higher for the sc- models than for the sc+ models (signed-rank test for congruent trials evaluated at 0.5SD noise: p = 0.0038; incongruent trials: p = 6.0e-4; stay trials: p = 5.5e-4; switch trials: p = 0.015; N = 25 models). Thus, the more compact latent representation of the sc- models resulted in a general degradation in performance, consistent with the view that noise in these models could more readily drive the latent state into incorrect response regions.

## Discussion

Many cognitive tasks—both inside and outside of the laboratory—require processing and responding to sequences of stimuli dynamically in real time. How can we identify models of the underlying cognitive and neural dynamics that remain faithful to this generative process? Most existing models fall short in that they either do not represent time at a fine enough timescale to model RT data, do not model multiple trials, or do not have the capacity to model individual differences in behavior. Leveraging large-scale cognitive training data and recent innovations in machine learning (*12, 22*), we developed a novel modeling framework—task-DyVA—in which expressive dynamical systems are trained to reproduce sequential behavior and RT data gathered from individual participants. We were able to recover interpretable dynamical models that precisely recapitulated subject-specific RT effects, making a compelling case for the fruitfulness of our approach.

Our approach joins a burgeoning research program in which deep neural network models are deployed to investigate neural and behavioral data (*6, 10, 23, 24*). Optimizing neural network models to perform cognitive tasks is an increasingly popular approach, one that has furnished compelling insights into the neural representation of behaviors as diverse as visual object recognition, context-dependent behavior, and flexible timing (*5–7*). However, a general conceptual issue with this approach is that human behavior often deviates systematically from optimality (*25, 26*), idiosyncrasies that a task-optimized neural network may fail to capture. Moreover, task-optimized networks cannot be readily extended to model individual differences in behavior. Like other recent work (*10, 23*), we address these limitations by using behavioral data to constrain the structure of our model. Unlike these previous methods, however, our model processes incoming stimulus information dynamically with sub-second temporal precision, allowing us to model sequences of RTs.

Our work provides support for a recently proposed normative account of the switch cost (*17*). Most prior work focuses on *how* the switch cost arises. Two prevailing viewpoints explain switch costs in terms of task representations or “task sets”: one view posits that cognitive control mechanisms require time to make task sets functionally active (*14*); the other posits that these task sets actively interfere with each other (*15*). In contrast, Musslick and Cohen provide an account of *why* the switch cost would exist in the first place (*17*). Building on prior models proposed by the authors themselves and others (*27–29*), they propose that the representation for each task is assigned to a unit whose time-varying activity corresponds to the degree of activation of the associated task. The units are self-exciting and mutually inhibitory, giving rise to two stable attractors, one for each task. Deeper attractors confer greater stability and resistance to distraction, but this comes at the expense of reduced behavioral flexibility: it takes more time to escape the attractor when a task switch is required, resulting in a switch cost.

Similarly, in our model, each task is associated with a separable set of stable fixed points within the latent space, forming two attracting regions. We demonstrate that the separation of the two task representations confers greater stability: models with less separated task representations were more affected by sensory noise. A notable difference between our work and existing attractor models of task-switching (*27–29*) is that we did not assume any particular functional relationship between the two task representations—e.g. by assigning each task to units that directly inhibit each other—or indeed that the two tasks should have a designated representation at all. The structured representation of the task that we observed arose emergently from the model architecture, objective function, and learning rules (*30*). Moreover, our model provides a means of directly testing the hypothesized relationship between neural representations and switch costs. In human or animal subjects performing a task-switching task with concurrent measurements of neural activity, one could quantify the separation of task representations within neural state space, much as we do here. If the performance of subjects with more separated task representations was less affected by sensory noise, this would support the notion that more separated task representations confer robustness.

While we chose to focus on a single well-studied task, allowing us to conduct a detailed study of the models’ learned task representation, the task-DyVA framework could be extended to model any cognitive task in which RTs are measured, in both human and animal subjects. More generally, our framework could also be adapted to incorporate other time-varying signals, including eye movements, cursor movements, and measurements of neural activity (*31*). In short, our framework provides a foundation for deriving generative models that explain how the brain dynamically gives rise to behavior.

## Acknowledgements

We thank J. Cunningham, G. Huckins, M. Steyvers, D. Sussillo, L. Tian, the Lumos Labs research team, and other members of the Poldrack Lab for helpful conversations and comments on the manuscript. We also thank Monico Chavez for providing graphics files of the Ebb and Flow stimuli. The authors acknowledge the Texas Advanced Computing Center (TACC) at The University of Texas at Austin for providing HPC and storage resources that contributed to the research results reported within this paper. URL: http://www.tacc.utexas.edu/.

## Data and materials availability

All code and data are publicly available without restriction. Trained models and data: https://doi.org/10.5281/zenodo.6368413. Code: https://github.com/pauljaffe/task-dyva/tree/v1.0.0.

## Author contributions

Conceptualization: PIJ, RAP, RJS, PGB; Methodology: PIJ; Software: PIJ; Validation: PIJ; Formal analysis: PIJ; Investigation: PIJ; Resources: RAP, RJS; Data curation: PIJ; Writing - original draft: PIJ; Writing - review and editing: PIJ, RAP, RJS, PGB; Visualization: PIJ; Supervision: RAP, RJS, PGB; Project administration: PIJ, RAP, RJS, PGB; Funding acquisition: RAP, RJS

## Supplementary Materials for

### Materials and Methods

#### Description of Ebb and Flow

Ebb and Flow is a game offered on the Lumosity cognitive training platform in which participants must switch between two cognitive tasks. On each trial, participants are shown a set of leaves that vary along two stimulus dimensions corresponding to the two tasks: the direction in which the leaves are pointing (left, right, up, or down) and the direction in which the leaves are moving (left, right, up, or down). Depending on the color of the leaves, participants must either report the pointing direction of the leaves or the direction of motion of the leaves with a key press (using the left, right, up, or down arrow keys). The pointing task is cued by green leaves while the motion task is cued by orange leaves. The cued task is also reinforced by verbal cues at the bottom of the screen (see Fig. 1A). After making a response, the participant is provided with brief visual feedback indicating whether or not the correct response was provided concurrent with the start of the next trial (correct: green check mark; incorrect: red/orange ‘X’ mark; both displayed briefly in the center of the screen). At the conclusion of a given gameplay event (duration = 60s), the participant is shown the number of correct trials out of the total number of attempted trials, the mean response time (RT), and a composite score based on the number of correct responses. Thus, the composite score is determined by a combination of response speed and accuracy.

When the pointing direction and motion direction of the leaves is the same, a trial is said to be congruent, and incongruent otherwise. The probability that any given trial will be congruent is 50%. The probability that the task cue will switch on a given trial is given by 0.05 × the number of trials since the last task switch. As such, the majority of trials for a given gameplay event are typically stay trials.

#### The task-DyVA representation of Ebb and Flow

##### Stimuli and response target

Participant data from Ebb and Flow was transformed into a simplified representation for use with the task-DyVA model. At each moment in time (step size = 20ms), a single binary-valued unit was used to represent the presence or absence of each stimulus modality × response direction and the two task cues (see Fig. 1B, bottom left). These stimulus inputs contribute directly to the dynamics of the latent state (see equation (1) below). For each trial, we allowed for the possibility that the model’s RT would be longer than the participant’s RT by extending the transformed stimuli 500ms beyond the participant’s RT. Zero-mean Gaussian noise (0.1 SD) was added independently to each stimulus unit. The noise was resampled on each iteration of the training loop. To create the response target used for model training, a Gaussian kernel (SD = 50ms, max = 1) was centered at each RT from the participant’s responses (see Fig. 1B, bottom right).

##### RT calculation

At the conclusion of training, the model typically generated time-localized activations at one of the four possible response directions on each trial (see Fig. 1B, bottom right). The direction of the model’s response on a given trial was calculated as the output unit with the maximum activation in a window beginning 100ms after stimulus onset and ending at stimulus offset. The RT for that trial was calculated as the temporal ‘center of mass’ of the output unit with the maximal activation:

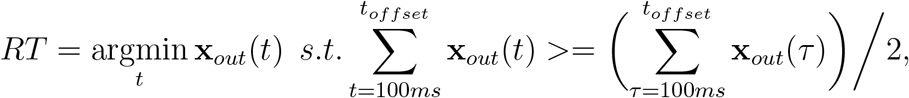

where **x**_*out*_(*t*) is the activation of the response unit at timestep *t*. RTs were not calculated for trials in which the stimuli were prematurely truncated.

#### The task-DyVA model

##### Generative model

The generative component of task-DyVA—i.e., the component that emulates human behavior—is a dynamical latent variable model that accepts sequences of task stimuli **u**_1:*T*_ as inputs and produces sequences of task responses **x**_1:*T*_ as outputs. Our implementation of the generative model is closely related to Deep Variational Bayes Filters (DVBF) (*22*), though the encoder model differs considerably (described below). The latent state variables **z**_1:*T*_ are defined to have locally-linear transition dynamics:

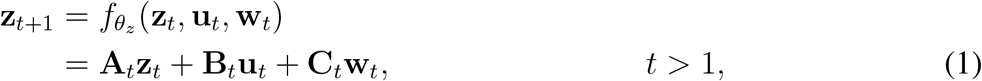

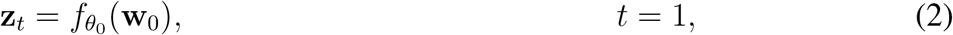

where the initialization function 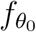 is parameterized by a MLP with two layers (hidden layer: 64 ReLU units; output layer: 16 linear units). When the model is used to generate task responses, i.e. at the conclusion of training, the stochastic variables **w**_*t*_ are sampled from the prior:

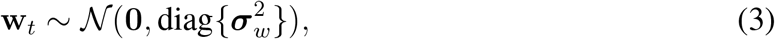

where the prior variance vector 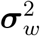 is learned. During model training, the **w**_*t*_ are instead sampled from the encoder model as described in the following section.

The matrices **A**_*t*_, **B**_*t*_, and **C**_*t*_ are linear combinations of time-independent matrices {**A**^(*i*)^, **B**^(*i*)^, **C**^(*i*)^}; *i* = 1, …, *M* that are learned as point estimates:

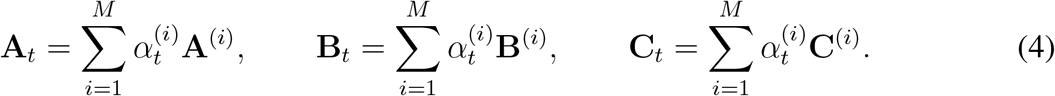

The weights *α*_*t*_ are the same for each set of matrices and are determined by a single-layer neural network 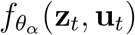 with softmax output units. For the models presented in this paper, **z**_*t*_ ∈ ℝ^16^, **w**_*t*_ ∈ ℝ^16^, **u**_*t*_ ∈ ℝ^10^, **x**_*t*_ ∈ ℝ^4^ and *M* = 2.

Model outputs **x**_*t*_, which map one-to-one onto each of the possible task response directions, are sampled from the decoder model according to:

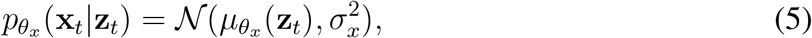

where 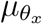 is parameterized by a MLP with two layers and learned parameters *θ*_*x*_ (hidden layer: 64 ReLU units; output layer: four sigmoid units). Note that for all analyses conducted on the trained models, e.g. for calculating RTs, we use 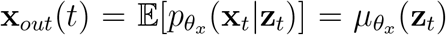 rather than stochastic samples. The decoder SD *σ*_*x*_, which is only relevant when evaluating the objective function, is fixed to 0.75. A probabilistic graphical model that depicts the dependencies of the variables in the generative model is shown in Fig. S1A.

##### Encoder model

To learn the parameters of the generative model *θ* = *θ*_*x*_ ∪ *θ*_*z*_, we make use of an approximate inference framework known as the variational autoencoder or VAE (*19, 20*). Approximate inference is essential for computationally efficient learning in our model since the conditional likelihood *p*_*θ*_(**x**_1:*T*_ |**u**_1:*T*_) is intractable due to nonlinearities in the dynamics equation (1) and decoder model. Following the VAE framework, we replace the true (intractable) posterior distribution in our model *p*_*θ*_(**w**_1:*T*_ |**u**_1:*T*_, **x**_1:*T*_) with a tractable approximate posterior distribution *q*_*ϕ*_(**w**_1:*T*_ |**u**_1:*T*_, **x**_1:*T*_) referred to as the encoder model that enables inference of the unobserved stochastic parameters **w**_1:*T*_ from the observed data {**x**_1:*T*_, **u**_1:*T*_}. Note that the posterior distribution is defined over the stochastic parameters **w**_1:*T*_ rather than the latent state variables **z**_1:*T*_ since the latent state evolves deterministically given **w**_1:*T*_ (*22*). The parameters of the encoder model *ϕ* are learned jointly alongside the generative model parameters *θ*. We note that the encoder model is only introduced to train the model and plays no role in generating responses once the model is fully trained.

The task-DyVA encoder model factorizes as follows:

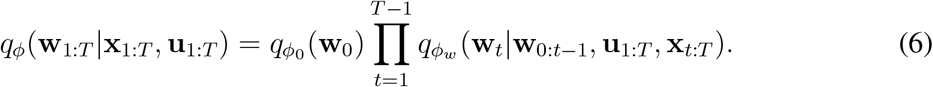

We chose this factorization since it is almost identical to the factorization of the exact posterior distribution, a strategy suggested by (*12*). Relative to the exact posterior distribution, the 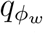 factors in our encoder model have additional dependencies on *u*_*T*_ and *x*_*t*_ (to be explicit: both refer to variables at a single timestep). These minor additional dependencies were incorporated for computational convenience. The exact posterior distribution can be derived using the chain rule of probability and the principle of D-separation applied to the graph of the generative model (Fig. S1A) (*32*).

To parameterize the density 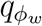, we introduce a backward RNN that transmits information from future observations to the current timestep. This information is encoded in the state variable 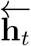. In the forward pass of the model, a neural network combines 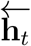 and **z**_*t*_ at each timestep, yielding parameters of a normal distribution ***µ***_*ϕ*_ and ***σ***_*ϕ*_ from which **w**_*t*_ is sampled. The dynamics of the latent state variable **z**_*t*_ are the same as those of the generative model defined in equations 1-5 (except that **w**_*t*_ is no longer sampled from the prior). Thus the encoder model is given by:

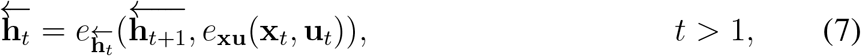

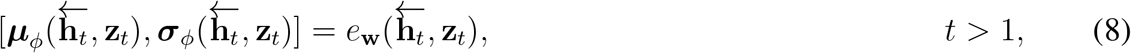

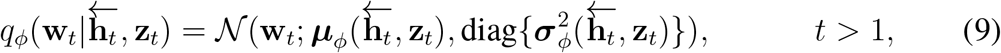

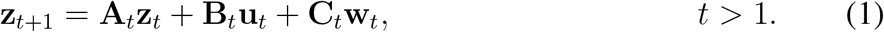

In equation (7), *e*_**xu**_ is parameterized by a MLP with two layers (hidden layer: 64 ReLU units; output layer: 64 linear units), and 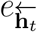 is parameterized by a single-layer LSTM RNN with 64 hidden units. In equation (8), ***µ***_*ϕ*_ and ***σ***_*ϕ*_ are both parameterized by a MLP with two layers and shared parameters in the hidden layer (hidden layer: 64 ReLU units). The output layer of ***µ***_*ϕ*_ consists of 16 linear units while the output layer of ***σ***_*ϕ*_ consists of 16 softmax units. After the softmax nonlinearity, the variance parameters are multiplied by a fixed factor of 16 and added to a small positive constant (1e-6). To initialize the model, **w**_0_ is sampled from the prior according to equation (3), so that 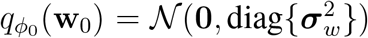, and **z**_1_ is given deterministically from the initialization network 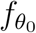 (which depends only on **w**_0_). A probabilistic graphical model showing the dependencies between the variables in the encoder model is shown in Fig. S1B.

##### Objective function

To train the task-DyVA model, we use a slightly modified form of the standard evidence lower bound (ELBO) objective function, so-called because it is a lower bound on the marginal likelihood. To calculate the ELBO, it will be helpful to derive the conditional density *p*_*θ*_(**x**_1:*T*_, **w**_1:*T*_ |**u**_1:*T*_). In what follows, we show how this density can be derived by marginalizing out the deterministic latent state variables **z**_1:*T*_ from the complete joint density of the generative model. An analogous calculation for a related model is provided in Appendix A of (*12*). The joint distribution of the latent and observed variables conditioned on the task stimuli is given by:

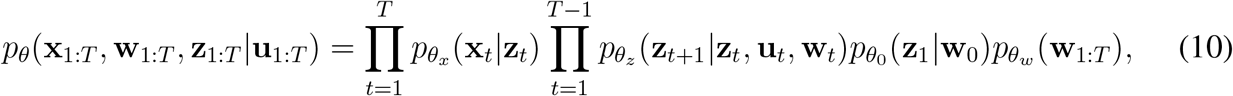

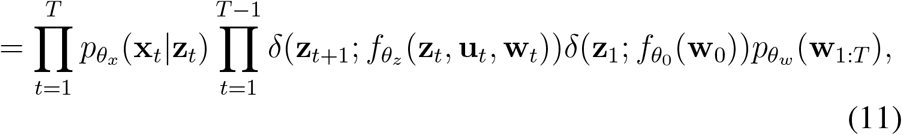

where the factorization on the right-hand side of equation (10) follows from the state space assumptions imposed on the generative model. The Dirac distributions in equation (11) result from the fact that **z**_*t*+1_ is a deterministic function of **z**_*t*_, **u**_*t*_, and **w**_*t*_ as specified in equations (1) and (2).

To marginalize over **z**_1:*T*_, we first express **z**_*t*_ as a deterministic function of **u**_1:*t*−1_ and **w**_1:*t*−1_ that results from the sequential application of 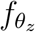 at each timestep. We denote the resulting expression as **d**_*t*_ so that:

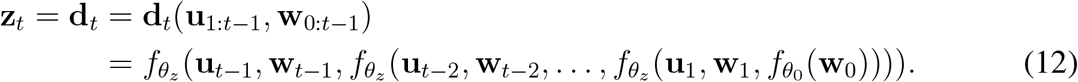

Inserting **d**_*t*_ into equation (11), and integrating out the **z**_1:*T*_ sequentially starting with **z**_*T*_, we derive the density:

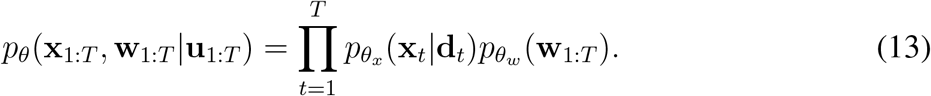

The ELBO objective function used to train task-DyVA is given by:

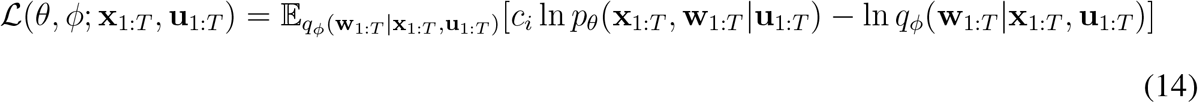

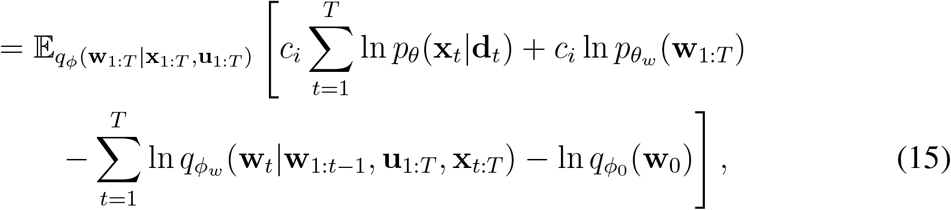

in which the densities defined in equations (6) and (13) have been inserted into equation (14). This is the usual ELBO objective with the modification that the joint likelihood is scaled by an annealing parameter that increases monotonically with the number of gradient updates according to *c*_*i*_ = min (1, 0.01 + *i/τ*_*A*_). Similar annealing schemes have been used in prior work (*22, 33*). For all models, *τ*_*A*_ was set to 40,000.

##### Related models

Due to the temporal dependency of our model, task-DyVA is a member of the recently characterized dynamical VAE family (*12*), from which we derive the name. Among prior work, the generative component of task-DyVA is most closely related to Deep Variational Bayes Filters (DVBF) (*22*). A significant point of departure between task-DyVA and DVBF is the encoder model: in task-DyVA, **w**_*t*_ depends on both the current and all future observations, which more closely resembles the structure of the true posterior distribution, while in DVBF only the observations **x**_*t*+1_ and **u**_*t*_ contribute to **w**_*t*_. The structure of the task-DyVA encoder model is perhaps most similar to a recent implementation of a recurrent VAE model (*34*).

#### Datasets

The analyses presented in this paper consist of models trained on deidentified data from 20 Lumosity participants in each of seven decade-long age bins (140 participants total; first age bin: ages 20 to 29, last age bin: ages 80 to 89). No statistical methods were used to predetermine sample sizes. These participants were randomly selected from a larger pool of participants who had played Ebb and Flow at least 500 times, who signed up as Lumosity participants between Jan. 1, 2012 and Dec. 31, 2021 (inclusive), who were between the ages of 20 and 89 at the time of signup (inclusive), whose country of origin was the United States, Canada, Australia, or New Zealand, who indicated that their preferred language was English, and who were not employees at Lumos Labs, Inc. or participants who created accounts for research purposes. We also required that each participant had only played Ebb and Flow on the web platform (as opposed to the mobile platform), that they did not pause for more than 100 days between consecutive plays of Ebb and Flow, and that their mean RT did not exceed 1.25s. Gameplays in which any RT exceeded 10s were not included in the datasets and not counted as valid gameplays (described below). Additional quality control checks were conducted to ensure that all game metadata was correctly coded. To be included in the pool of available participants, a given participant had to have at least 300 valid gameplays between their 150th and 500th gameplay (inclusive). The final data from a given participant consisted of all valid gameplays between that participant’s 150th and 500th gameplay (inclusive). Additional preprocessing steps are described in the model training section.

In this set of 140 participants, 46% identified as female, 44% identified as male, and 10% did not report their gender. These participants reported their educational attainment as follows: some high school (1%), high school (15%), some college (19%), college degree (21%), associate’s degree (7%), master’s degree (15%), professional degree (7%), Ph.D. (2%), other (3%), with 9% not reporting education information. In an additional small subset of participants, the training loss diverged to infinity; these failed models are not included in summary analyses (20 additional participants corresponding to a failure rate of 12.5%).

Prior to training models on these 140 participants, we used a different set of ‘exploratory’ participants randomly selected from the available pool to develop the task-DyVA modeling framework (five to ten participants in each age bin). All model parameters were established on this exploratory subset; models trained on these participants were not included in summary analyses. This was done to provide some assurance that the model parameters presented in this study would work well if applied to data from other participants.

#### Model training

##### Training, validation, and test sets

Gameplay data from each participant was randomly divided into three splits: a training set (50%), a validation set (20%), and a holdout/test set (30%). Following standard procedure, the validation set was used to monitor the progression of training and was not used to update the model’s parameters. The test set was used to assess the model’s ability to generalize to unseen data; all summary analyses are reported for model responses generated on the test set.

##### Data augmentation

A data augmentation approach was adopted to increase the quantity of training data and to balance the proportion of stay and switch trials in the training set. Broadly, this was accomplished by composing training data as a mixture of the original trial sequences and sequences resampled from estimated RT PDFs. Specifically, we segmented the original 60s-long gameplay data into short sequences (5s duration, 3-4 trials), where the first sequence began at 5s and the last sequence began at 55s. Using the RTs from these trial sequences, we estimated four RT PDFs corresponding to four different types of trials: congruent stay trials, congruent switch trials, incongruent stay trials, and incongruent switch trials. Each PDF was estimated as a Gaussian kernel-density estimate (KDE; scale parameter = 0.25). Prior to estimating these PDFs, we excluded trial sequences with extreme outlier RTs as described in a subsequent section.

To increase the quantity of training data, the corpus of 5s trial sequences was replicated by a factor of ten (yielding ten total copies of the original corpus). Then, for each trial sequence, we randomly determined if that sequence would be used as is (25% of the time), or would be replaced with a synthetic trial sequence resampled from the RT KDE (75% of the time). In the latter case, we first randomly sampled one of the four KDEs with equal probability (i.e. congruent/stay, congruent/switch, incongruent/stay, or incongruent/switch), then randomly sampled a RT from the corresponding KDE. The stimuli for that trial were sampled randomly with equal probability from the set of permissible stimuli that were determined by the sampled trial type (e.g. congruent vs. incongruent); the task cue was determined implicitly by the sampled trial type. The response direction for that trial was set to be the correct response direction given the stimuli and task cue. This resampling process was repeated until the cumulative duration of the trials in the sequence exceeded 5s. Finally, the trial sequence was transformed to a continuous representation as described in a previous section.

The resampling procedure described above was done only for the training set: the validation and test sets used the original trial sequences as observed in the data. For the validation and test sets, the original gameplay data were segmented into 10s-long sequences (rather than 5s-long sequences), where the first sequence began at 5s and the last sequence began at 55s. For all three splits (training, validation, and test), trial sequences with outlier RTs were excluded (described below), as were trial sequences that did not contain at least one complete trial.

For the models trained on synthetic data with no switch cost (the scmodels), the data augmentation procedure described above was modified to eliminate the switch cost. As before, we determined RT KDEs for each of four different trial types: (congruent/stay, congruent/switch, incongruent/stay, and incongruent/switch). To eliminate the switch cost on congruent trials, we translated the congruent/switch KDE along the time axis such that the mean was the same as that of the congruent/stay KDE. Similarly, the incongruent/switch KDE was translated to eliminate the switch cost on incongruent trials. Finally, all four KDEs were translated by the same amount, such that the mean RT of the KDEs with no switch cost would be the same as that of the mean RT of the original KDEs (where the mean RT is calculated by sampling from each of the four KDEs with equal probability). Additionally, instead of constructing synthetic data for 75% of trial sequences, where the remaining 25% of sequences were unmodified, all trial sequences were resampled. All other aspects of the modified data augmentation procedure were the same as described above.

##### Outlier removal

Trial sequences in which any RT was categorized as an outlier were excluded. The outlier procedure made use of the median absolute deviation from the median (the MAD). An RT was classified as an outlier if the absolute deviation from the median exceeded ten times the MAD from that participant’s data. Thus, only extremely short and extremely long RTs were classified as outliers. The MAD and median were estimated from all RTs in the participant’s data.

##### Early stopping

To guard against overfitting, we monitored the progression of model training using the validation set and halted training after certain criteria were met. The metric we monitored, *m*_*stop*_, depended on the difference between model and participant behavior (in particular, the switch cost and congruency effect):

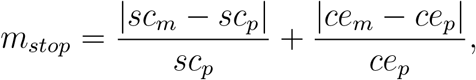

where *sc*_*m*_ and *sc*_*p*_ are the model and participant switch cost, respectively, and *ce*_*m*_ and *ce*_*p*_ are the model and participant congruency effect (all metrics were evaluated on the validation set).

*m*_*stop*_ was calculated every 10th training epoch, beginning with the 500th epoch. Training was halted if *m*_*stop*_ did not decrease for 200 consecutive epochs. The training epoch with the lowest value of *m*_*stop*_—as evaluated on the validation set—was used for all model analyses. We reiterate that the model responses presented in the main text were generated using a holdout test set (separate from the validation set).

A modified early stopping criterion was used for the models trained on synthetic data with no switch cost (the sc- models), since the goal of model training was to eliminate the switch cost rather than approximate the participant’s switch cost. Specifically, we used the simple stopping criterion 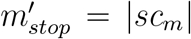, where *sc*_*m*_ is the model switch cost. As described above, the training epoch with the lowest value of 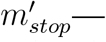evaluated on the validation set—was used for model analyses. All other aspects of the modified early stopping procedure were the same as described above.

##### Training models with no switch cost

We trained 25 models corresponding to 25 participants using synthetic data constructed to have no switch cost (the sc- models). To enable paired comparisons, these participants were selected from the main pool of 140 participants. Specifically, we rank-ordered all 140 participants by their switch cost, then trained models sequentially starting with the participant with the highest switch cost until 25 models were successfully trained. It was not possible to predetermine the exact set of participants that would be used for these experiments since the training loss sometimes diverged to infinity; in these cases, the model training was considered to be unsuccessful, and another participant was selected. The training failure rate in this set of models was somewhat higher than that observed in the original set of 140 models (37.5% vs. 12.5%). Other relevant details are described in the last paragraphs of the data augmentation and early stopping sections. Unless noted otherwise, all other aspects of model training and analysis were the same as described for the main set of 140 models.

##### Hyperparameters and other details of model training

The same model architecture and hyper-parameters were used to train all models, and all models were initialized with the same random seed. All models were trained on a NVIDIA GTX 1080 Ti GPU and/or a NVIDIA Tesla P100 GPU.

We used the AMSGrad variant of the Adam algorithm (*35*) as implemented in PyTorch version 1.8.1. The following Adam parameters were used for all model training: learning rate = 1e-4, *β*_1_ = 0.99, and *β*_2_ = 0.999. The training batch size was 128.

To avoid pathologically large gradients, we used gradient clipping with a clipping value of five: when the Euclidean norm of the parameter vector exceeded five, gradients were rescaled to have a norm of five. The norm was computed over all parameters concatenated into a single vector.

#### Model analysis methods

##### Overview

Unless noted otherwise, all summary analyses were performed on the holdout/test dataset, and analyses of the latent state variables were conducted in the full 16-dimensional latent space.

##### PCA

To visualize the latent state trajectories, the latent variables were projected onto the top three PCs determined from that model’s latent state matrix. The latent state matrix for a given model was formed by concatenating all of latent state sequences obtained from the test set along the time axis.

##### LDA

For the LDA analyses presented in Fig. 3C, latent state trajectories for all trials were aligned to stimulus onset and categorized according to the task cue and response direction. The positions of the latent state trajectories evaluated at the model’s mean RT were used to train the LDA models. Classification was performed using the full 16-dimensional latent state vectors. Since our goal was to use these LDA models to quantify the separability of the latent state vectors in each class, rather than make out-of-sample predictions, misclassification rates were calculated on the same data that was used to train the models. For the response direction classification models, we trained a separate LDA model for each combination of task cue x response direction, then averaged the misclassification rates across these eight models to obtain a single value for each participant’s model. To construct the shuffled datasets, we pooled data from both classes, then randomly reassigned the class labels. For the task classification models, we trained a separate LDA model on each of 100 shuffled datasets, then averaged the misclassification rates across these 100 models. For the response direction classification models, we calculated the average shuffle misclassification rate on each of 100 shuffled datasets for each task cue x response direction, then averaged the shuffled error rates to obtain a single value for each participant’s model.

##### Fixed points

Stable fixed points were identified by running the models in ‘generative mode’ using long (50s) sequences of static stimuli as inputs. If the model latents converged to a stable state, the point to which the latent state converged was identified as a stable fixed point. Specifically, for a given latent state sequence, we calculated the s.d. of each latent variable over the last 10s of the sequence. If the mean SD—averaged across the latent variables—was less than or equal to 0.001, we defined the time-averaged latent state variables over the last 10s of the sequence as a stable fixed point. For each model, we screened for fixed points using each of the 32 possible stimulus configurations, initializing the model from 10 different random locations for each stimulus configuration. These initial states were randomly sampled from all timepoints and all trials visited by the latent state from the responses generated from the test set. For a given stimulus configuration, the latent state sequences generated from the 10 initial states almost always converged to the same fixed point. However, if the Euclidean distance of the end state of a given latent state sequence was greater than 0.001 from all other fixed points identified with that stimulus configuration, the end state was identified as a new fixed point.

We identified stable fixed points associated with both task cues in 137 out of 140 models. Of the remaining three models, one did not have any stable fixed points, and the other two only had fixed points for one of the task cues. These three models were excluded from the analyses in which we calculated distances between pairs of fixed points. For the 137 models included in these analyses, the vast majority of models (135 out of 137) had at least 10 fixed points (mean ± s.e.m. number of fixed points per model: 25.8 ± 0.55). Averaged across these 137 models, 80 ± 1.7% (mean ± s.e.m.) of the 32 possible stimulus configurations were associated with at least one stable fixed point. In some cases, we observed oscillatory rather than static limiting behavior (i.e. stable limit cycles); these cases were not analyzed further.

##### Distance measures and task centroids

The task centroid for a given task was defined as the centroid of the latent state trajectories evaluated at stimulus onset for all stay trials in which the corresponding task cue was active (latent state trajectories were aligned to stimulus onset). To measure the distance between fixed points (Fig. 3D) and the distance between the latent state at trial onset and task centroids (Fig. 4B), we used the Euclidean distance calculated in the full 16-dimensional latent space. To calculate the distance between task centroids (Figs. 4C and 4E), we used a normalized distance measure that accounts for potential differences in the overall scale of the latent state variables across different models. This normalized distance *D*_*N*_ was defined as the Euclidean distance *D*_*E*_ divided by a scaled volume measure of the latent state variables from that model:

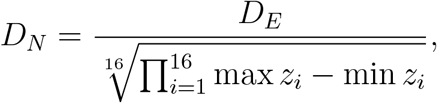

where the *z*_*i*_ denote the latent state variables, and the maxima and minima are calculated over all trials and all timepoints in the responses generated on the test set.

For the analyses in which we calculated the Pearson’s *r* between the distance to the task centroid and RTs on switch trials (Fig. 4B), *r* was calculated separately for trials with a fixed stimulus configuration, then averaged across stimulus configurations to obtain a single correlation estimate for each model. A stimulus configuration is defined by the values of the moving stimuli, pointing stimuli, and task cue on the current trial, yielding 32 possible stimulus configurations. We excluded stimulus configurations with fewer than five trials from the average Pearson’s *r* calculation (18 out of 140 models had at least one excluded stimulus configuration; mean ± s.e.m. excluded stimulus configurations within those models: 1.5 ± 0.16). One additional stimulus configuration from one model was excluded because the RTs were constant (thus, Pearson’s *r* is undefined).

#### Statistics

All reported statistics were calculated using two-sided tests. All assertions about statistical significance refer to a significance level *α* of 0.05. All p-values reported for signed-rank tests were derived using a normal approximation (rather than the exact null distribution of the test statistic). Additional details are provided throughout the text.

**Fig. S1:**
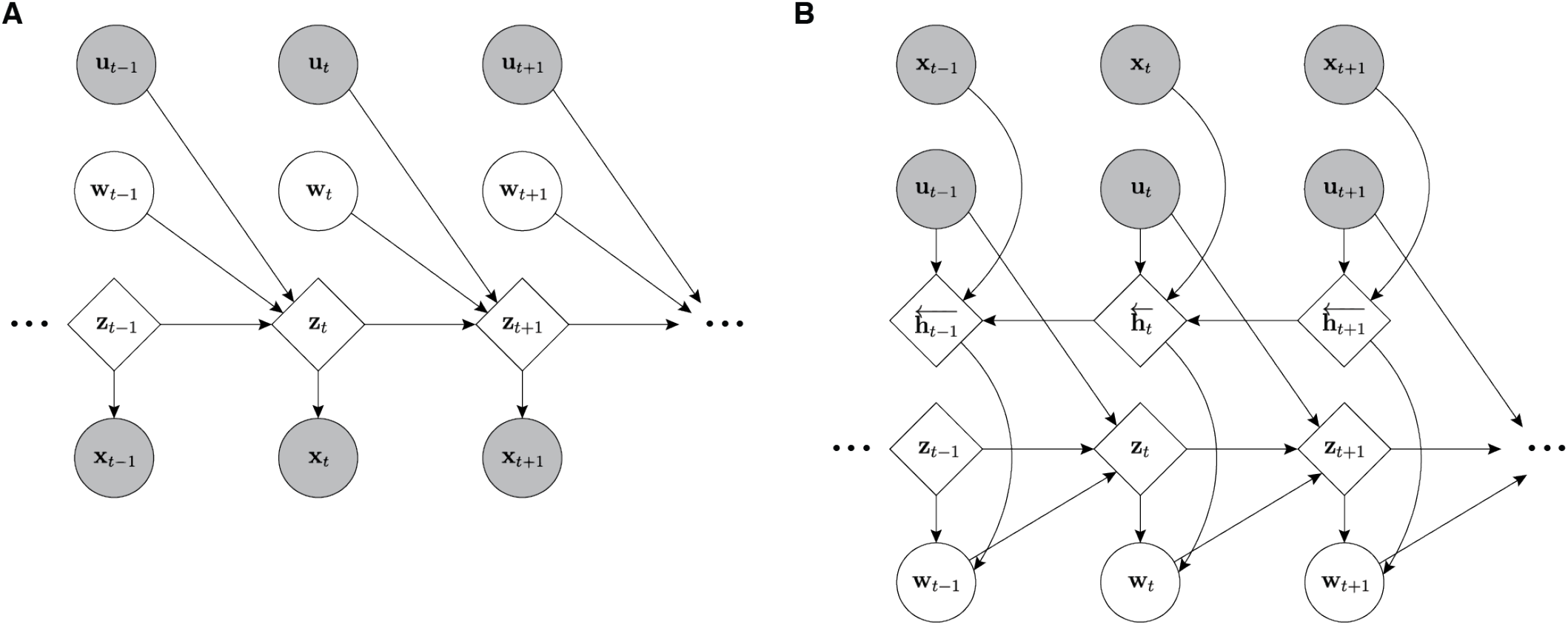
Probabilistic graphical models of task-DyVA. Shaded nodes indicate observed variables; unshaded nodes indicate unobserved (latent) variables. Circles indicate variables that depend stochastically on their parent nodes; diamonds indicate variables that depend deterministically on their parent nodes. See the Materials and Methods for a description of the variables. (**A**) The generative model of task-DyVA. (**B**) The encoder or inference model of task-DyVA.

**Fig. S2:**
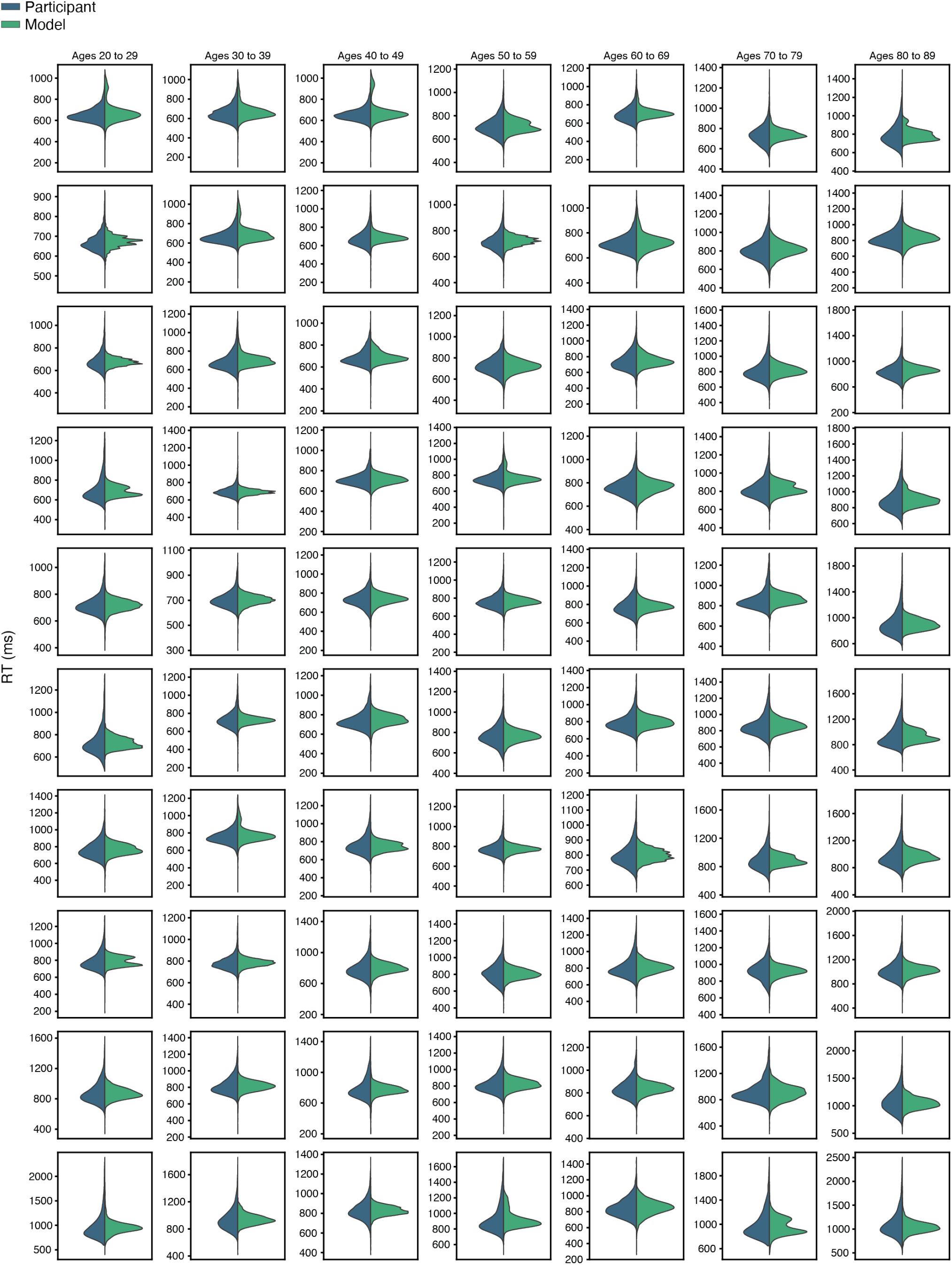
Example RT distributions from participants and fitted models. Within each age bin (columns), participants were sorted by their mean RT; every other participant is shown (10 out of 20 participants within each age bin; top row = shortest RTs).

**Fig. S3:**
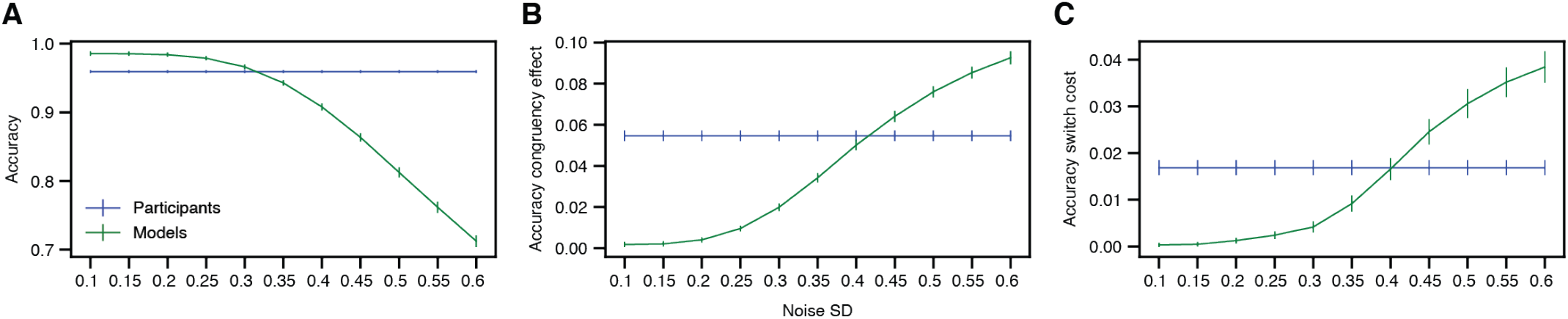
Increasing stimulus noise reveals effects on model accuracy. (**A**) Mean ± s.e.m. accuracy across models vs. stimulus noise SD (N = 140 participants/models). Note that participant accuracy (blue) was not assessed at different noise levels. At the noise level used to train the models (0.1SD), the mean ± s.e.m. accuracy was 0.99 ± 0.0033 for the models and 0.96 ± 0.0022 for the participants. (**B**) As in panel (A), but for the accuracy congruency effect (accuracy on congruent trials minus accuracy on incongruent trials). At 0.4SD noise, the mean ± s.e.m. accuracy congruency effect was 0.050 ± 0.0026 for the models and 0.055 ± 0.0026 for the participants (p = 0.14, signed-rank test). (**C**) As in panel (A), but for the accuracy switch cost (accuracy on stay trials minus accuracy on switch trials). At 0.4SD noise, the mean ± s.e.m. accuracy switch cost was 0.017 ± 0.0024 for the models and 0.017 ± 0.0016 for the participants (p = 0.97, signed-rank test).

**Fig. S4:**
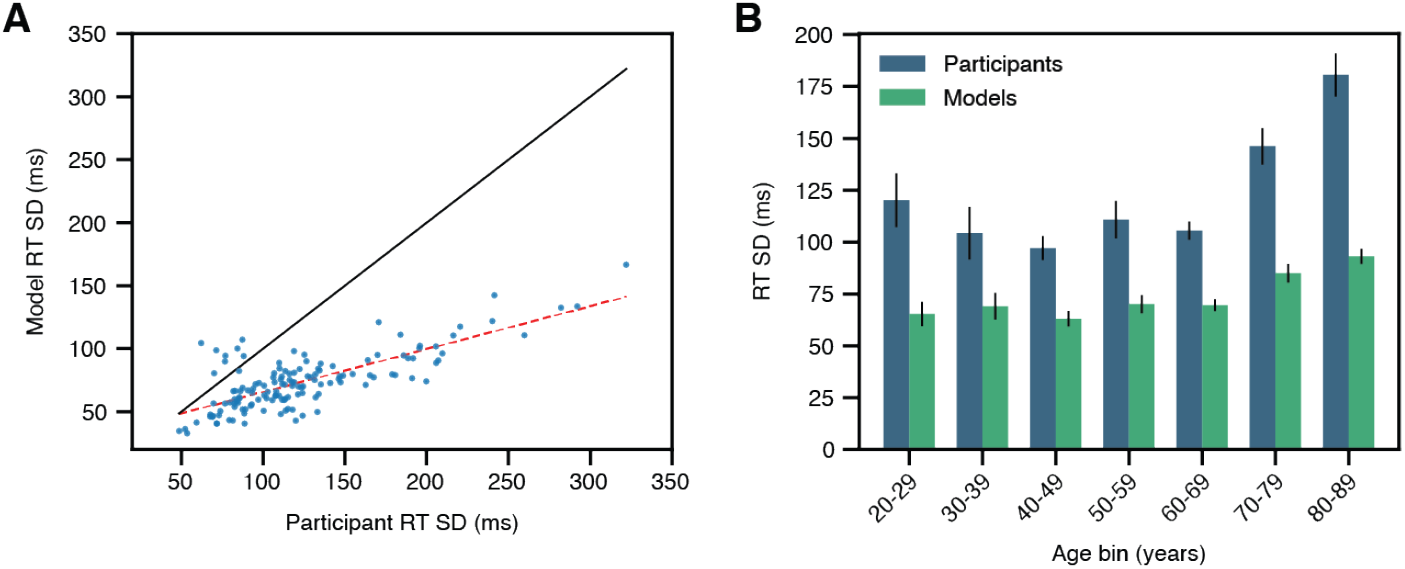
Comparison of model and participant RT variability. (**A**) RT standard deviation (SD) for participants and corresponding models (each point is one participant/model; black line: unity; red dashed line: best linear fit, slope = 0.34; Pearson’s *r* = 0.76, test for non-zero correlation using the exact distribution of *r*: p < 1e-26; N = 140 participants/models). The mean ± s.e.m. RT SD for the models was 73.6 ± 1.9ms and 123.5 ± 4.2ms for the participants. (**B**) Mean ± s.e.m. RT SD within each age bin (N = 20 participants/models per age bin).

**Fig. S5:**
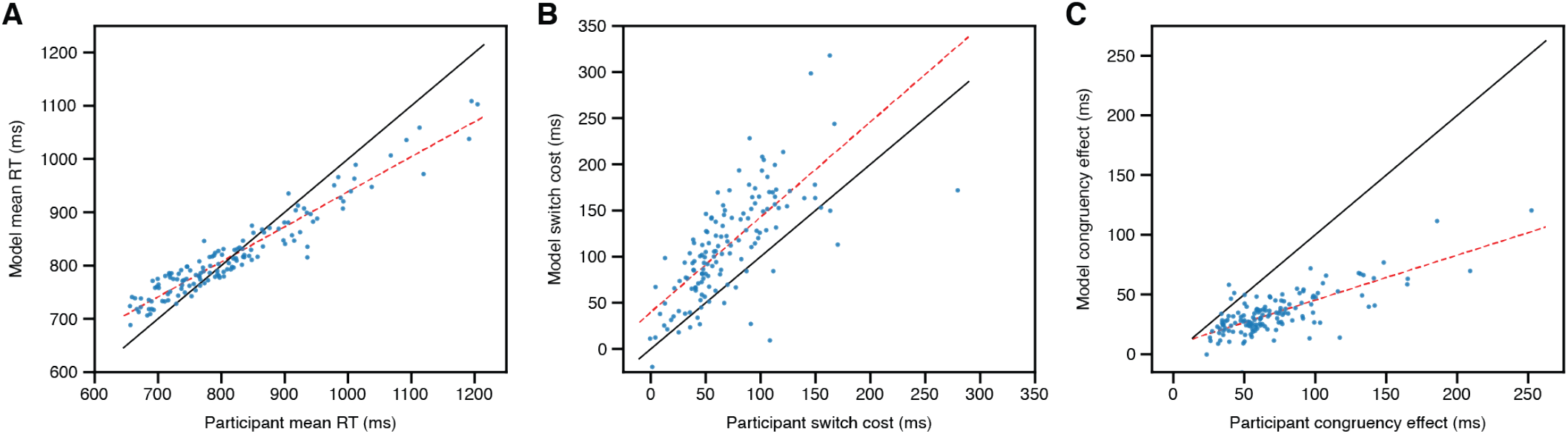
Strong correlations between participant and model behavioral metrics are preserved with elevated stimulus noise. (**A**) Mean RTs at 0.4SD stimulus noise (Pearson’s *r* for participants vs. models: 0.96, test for non-zero correlation using the exact distribution of *r*: p < 1e-75, best-fit slope = 0.66). (**B**) Switch costs at 0.4SD stimulus noise (Pearson’s *r* for participants vs. models: 0.70, p < 1e-21, best-fit slope = 1.0). (**C**) Congruency effects at 0.4SD stimulus noise (Pearson’s *r* for participants vs. models: 0.76, p < 1e-26, best-fit slope = 0.38). For panels A-C, each point is one participant/model; black line: unity; red dashed line: best linear fit; N = 140 participants/models. Note that participant behavior was not assessed at different noise levels.

**Fig. S6:**
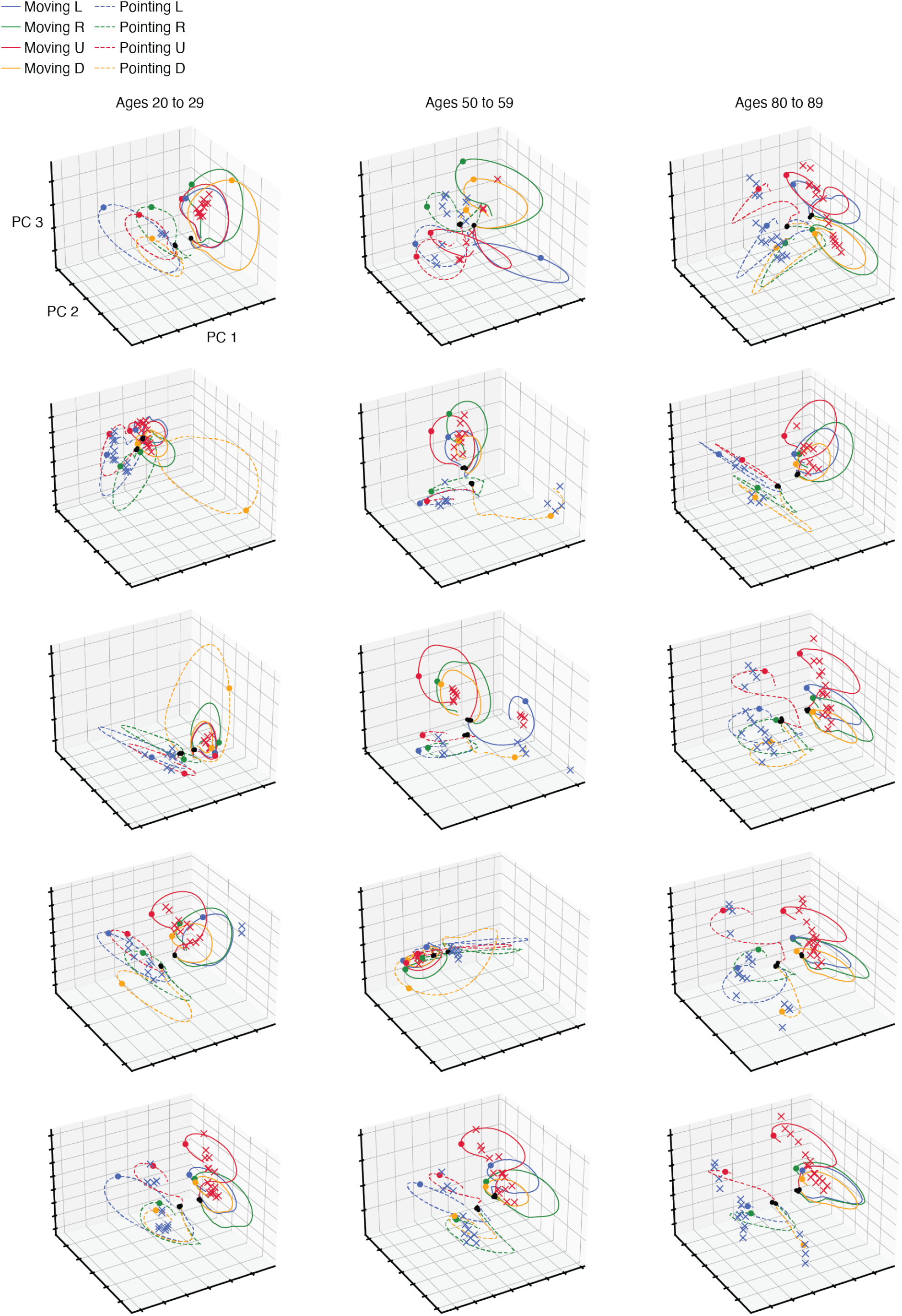
Latent representations of additional trained models. Each plot shows the trial-averaged latent state trajectories and stable fixed points (‘x’ marks) from one model. Within each age bin (columns), participants were sorted by their mean RT; every fourth participant is shown (5 out of 20 participants within each age bin; top row = shortest RTs).

**Fig. S7:**
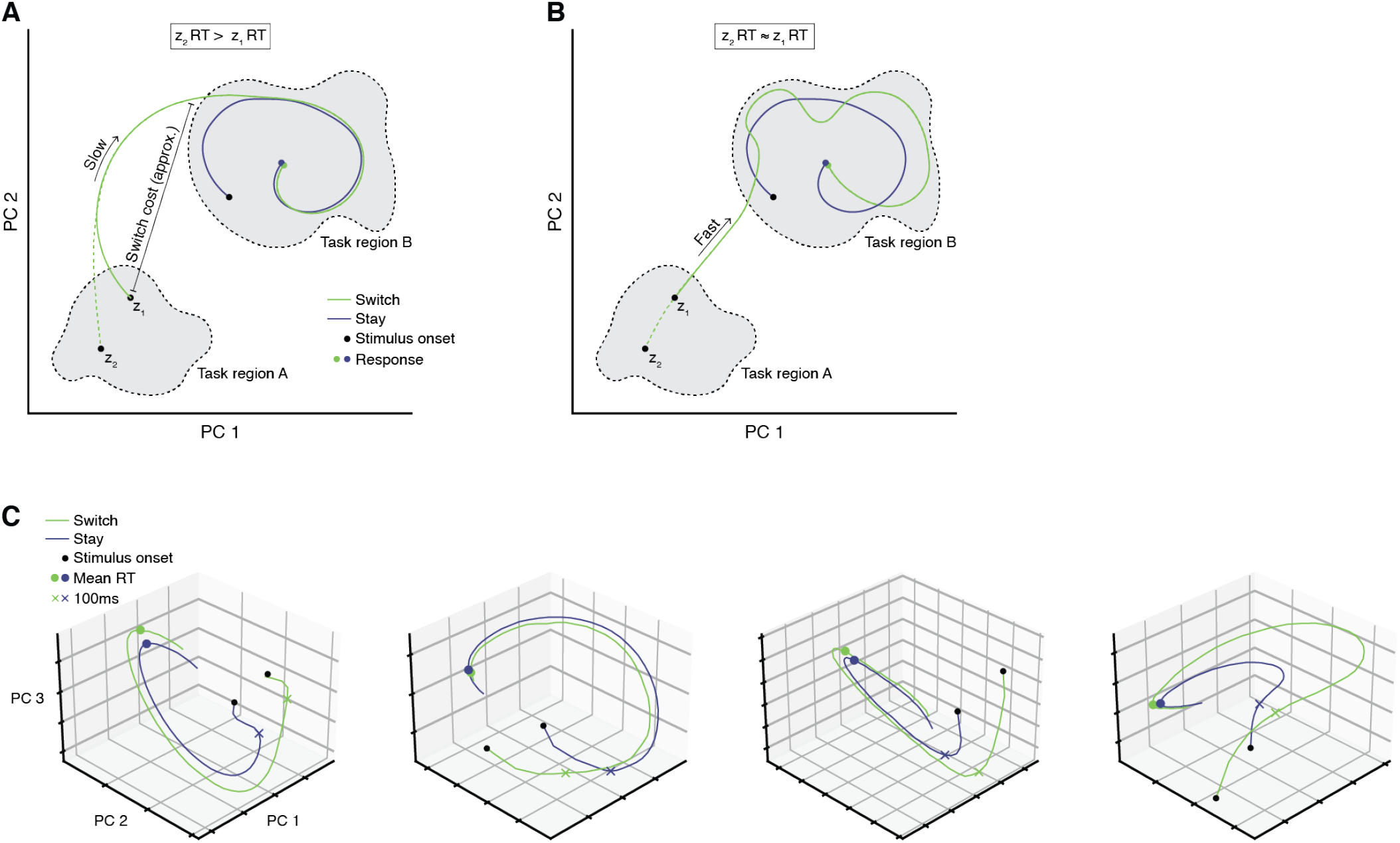
Hypothetical factors contributing to switch costs and example trajectories. (**A**) Illustration of how greater separation between task regions could contribute to switch costs. The transition from task region A to task region B is costly (i.e. slow). This model predicts a positive correlation between the distance to task region A at stimulus onset and RTs for switch trials. (**B**) Illustration of how slower dynamics within task region B could contribute to switch costs. The transition from task region A to task region B is fast. This model predicts that the correlation between the distance to task region A at stimulus onset and RTs for switch trials will be near zero. (**C**) Example latent state trajectories from four models. For each model, the stay and switch trajectories were averaged over trials with a fixed stimulus configuration (same stimuli and task cues on the current trial; task cues on the previous trial differed to select stay vs. switch trials). The stimulus configurations varied across the four models.

**Fig. S8:**
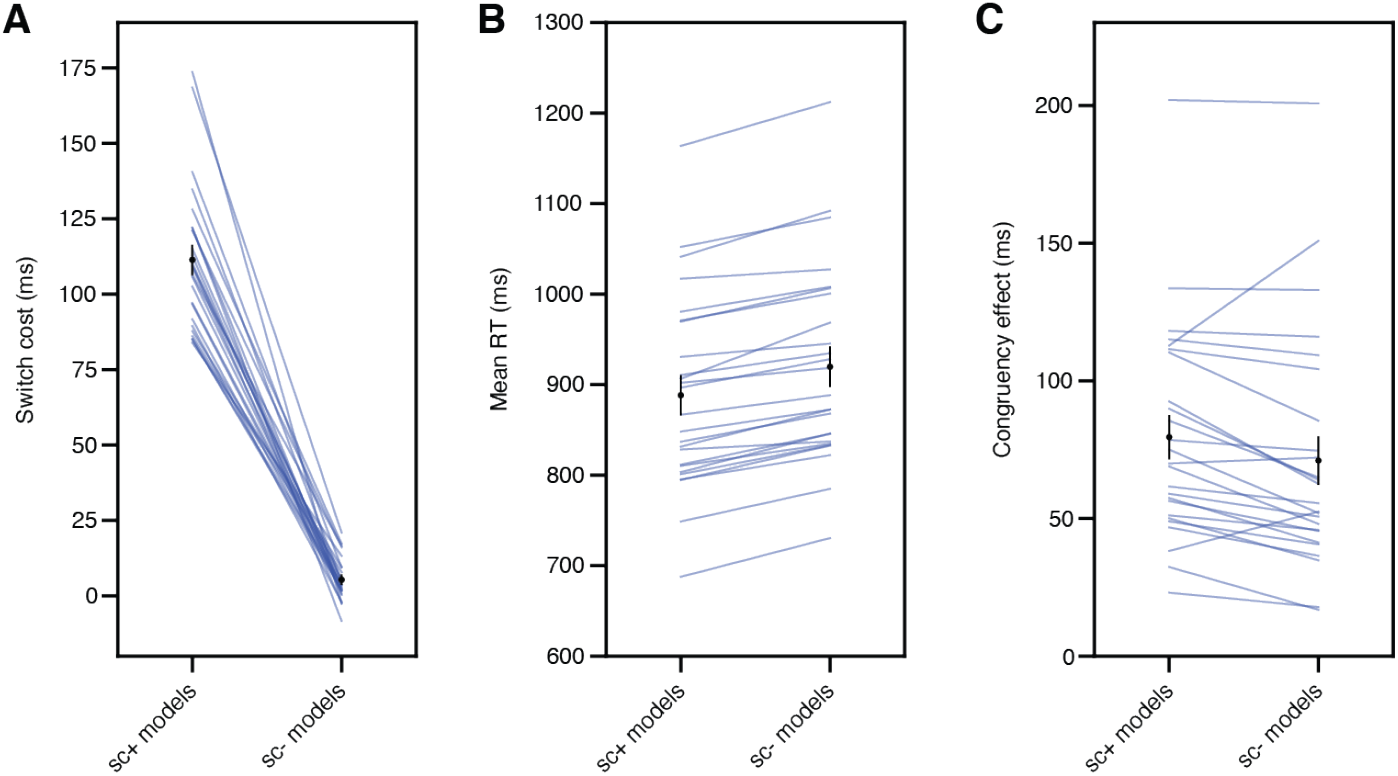
The sc- models exhibit reduced switch costs. (**A**) Switch cost for the sc+ and sc- models (black: mean ± s.e.m.; blue lines connect models trained on data from the same participant; mean ± s.e.m. for sc+ models: 111.4 ± 4.7ms; sc- models: 5.4 ± 1.4ms; N = 25 model pairs). (**B**) Mean RT for the sc+ and sc- models (mean ± s.e.m. for sc+ models: 888.4 ± 21.3ms; sc- models: 920.0 ± 21.6ms; N = 25 model pairs). (**C**) Congruency effect for the sc+ and sc- models (mean ± s.e.m. for sc+ models: 79.6 ± 7.7ms; sc- models: 71.1 ± 8.5ms; N = 25 model pairs).

**Fig. S9:**
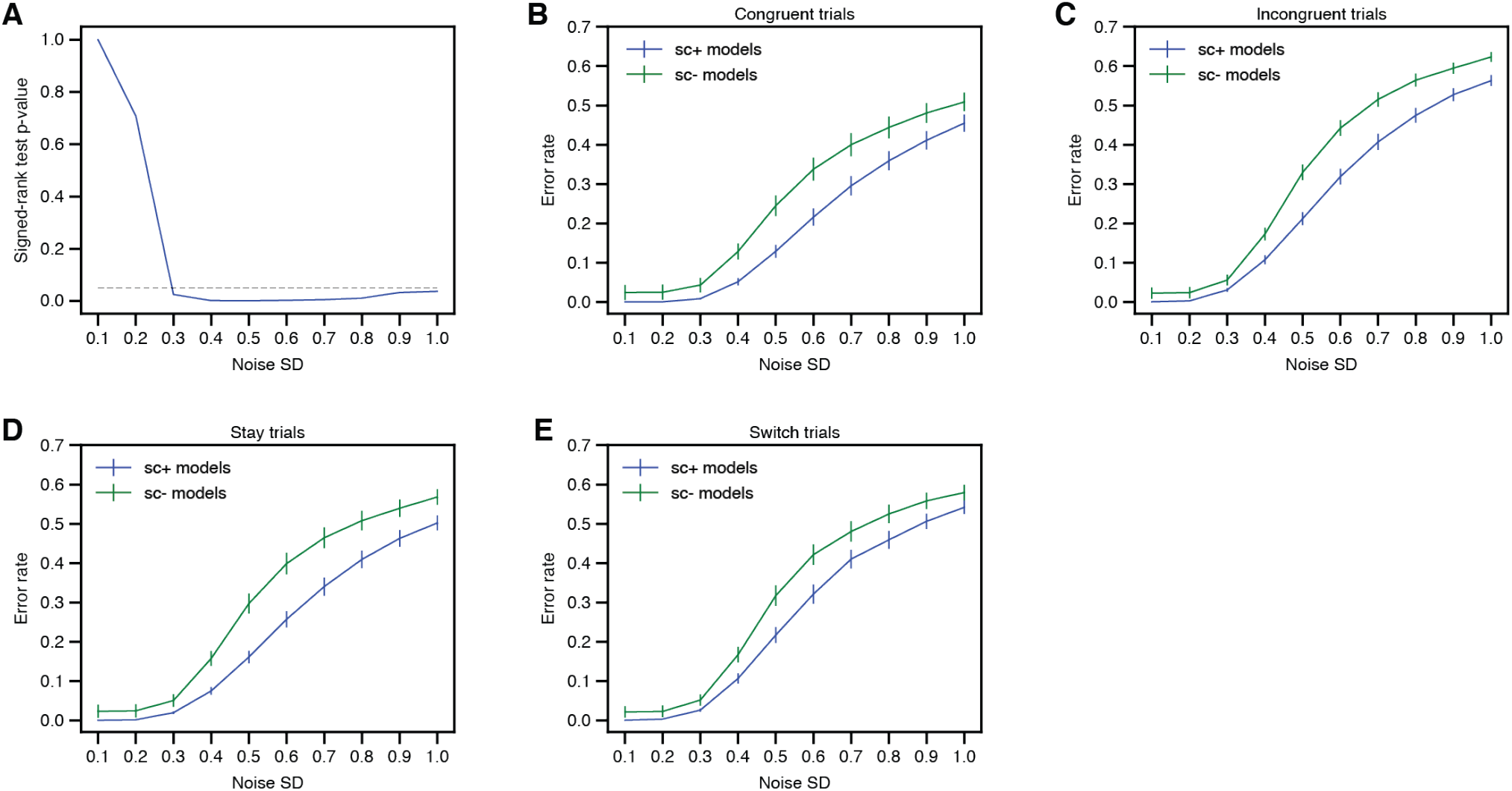
The reduced accuracy of the sc- models is consistent across noise levels and trial types. (**A**) Signed-rank test p-value for sc+ vs. sc- model accuracy (N = 25 model pairs; black dashed line: p = 0.05). (**B**) Mean ± s.e.m. error rate for congruent trials. (**C**) Mean ± s.e.m. error rate for incongruent trials. (**D**) Mean ± s.e.m. error rate for stay trials. (**E**) Mean ± s.e.m. error rate for switch trials. Panels B-E: N = 25 sc+ models and 25 sc- models.

